# A Novel Regulatory Role for the Circadian Clock Protein TOC1 via RNA binding

**DOI:** 10.1101/2020.05.06.071340

**Authors:** Yi Li, Yong-Gang Chang, Dawn H. Nagel, Tingjian Chen, Matias L Rugnone, Andy LiWang, Steve A. Kay

**Affiliations:** Department of Neurology, University of Southern California, Los Angeles, CA 90089, United States; School of Natural Sciences, University of California at Merced, Merced, CA 95343, United States; Chemistry & Chemical Biology, University of California at Merced, Merced, CA 95343, United States; Quantitative & Systems Biology, University of California at Merced, Merced, CA 95343, United States; Center for Circadian Biology, University of California at San Diego, La Jolla, CA 92093, United States; Health Sciences Research Institute, University of California at Merced, Merced, CA 95343, United States; The Center for Cellular & Biomolecular Machines, University of California at Merced, Merced, CA 95343, United States; Department of Botany and Plant Sciences, University of California in Riverside, Riverside, CA 92521, United States; School of Biology and Biological Engineering, South China University of Technology, Guangzhou, 510006, China; Monash Biomedicine Discovery Institute, Monash University, Clayton VIC, 3800, Australia

**Keywords:** circadian clock, transcriptional regulation, RNA binding protein

## Abstract

The circadian clock enables plants to predict daily changes of external signals and synchronize them with internal processes, conferring enhanced fitness and growth vigor. The first described *Arabidopsis* circadian clock protein is TIMING OF CAB EXPRESSION 1 (TOC1), which functions in a transcriptional feedback loop with two myb transcription factors, CIRCADIAN CLOCK-ASSOCIATED 1 (CCA1) and LATE ELONGATED HYPOCOTYL (LHY). Previous studies have shown that TOC1 is a DNA-binding transcriptional repressor of *CCA1* and *LHY*. However, the DNA motifs enriched amongst TOC1 targets share weak sequence similarity and lack consensus, suggesting that TOC1 regulates the expression of its targets through a novel mechanism. Here we show that the TOC1 protein binds directly to RNA via its conserved CCT domain. Using in vitro RNA selection, we identified an RNA motif that is recognized by the TOC1-CCT domain. The TOC1-CCT domain binds to this RNA sequence with nanomolar affinity determined by quantitative electrophoretic mobility shift assays (EMSAs) and isothermal titration calorimetry (ITC). NMR experiments showed that two CCT fragments, CCT^533-547^ and CCT^550-565^, use basic residues to bind the RNA motif. Mutational analysis confirmed that lysyl and arginyl residues bind to RNA in a cooperative manner. Furthermore, transiently expressed wildtype and mutant TOC1 in protoplasts demonstrated that RNA binding activity of TOC1 is required for its function as a transcriptional repressor in vivo. Our results reveal a novel regulatory mechanism for TOC1 through RNA binding, suggesting that TOC1 might play key roles as a multi-function protein.

## Introduction

As an adaptation to the external environment, an internal and self-sustaining timekeeping mechanism known as the circadian clock helps organisms to anticipate day-night changes so they can optimize the timing of resource allocation^1^, and modulate responses to occur under the most appropriate conditions^2,3^. In *Arabidopsis thaliana*, the core molecular components of this timekeeper consist of TIMING OF CAB EXPRESSION 1 (TOC1)^4,5^ along with two myb-like transcription factors CIRCADIAN CLOCK-ASSOCIATED 1 (CCA1)^6,7^ and LATE ELONGATED HYPOCOTYL (LHY)^8,9,10^ that function as the central oscillator. CCA1 and LHY are transcriptional repressors that directly bind to the evening element (EE)^11^, a DNA motif enriched in the promoters of many evening expressed genes including *TOC1*^12^.

More than 15 years ago, TOC1 was identified in a screen for mutants with shortened circadian period using a reporter line containing the promoter of the *LIGHT-HARVESTING CHLOROPHYLL a/b BINDING PROTEIN* gene (*LHCB1*.*1*; also called *CAB2*) fused to firefly *LUCIFERASE* (*LUC*) (*CAB2:LUC*)^4^. Extensive genetic studies showed that TOC1 is a positive component in the core clock, since reactivation of *CCA1* and *LHY* expression in the morning is dependent on TOC1^11^. However, overexpression of *TOC1* results in low mRNA levels of *CCA1* and *LHY*^5,13,14^, suggesting that TOC1 also acts as a transcriptional repressor. TOC1 belongs to the plant specific PSEUDO-RESPONSE REGULATORS (PRR) protein family^15^, whose five members all share three distinct domains: an amino-terminal PRR domain, an intermediate region (IR) and a carboxyl-terminal CCT (for CONSTANS, CONSTANS-LIKE, and TOC1) domain. The PRR domain shares high sequence similarity with the receiver domain of a two-component response regulator in bacteria^16^, but lacks the conserved aspartate residue that accepts a phosphoryl group from the sensor kinase. The PRR domain has also been shown to be involved in protein-protein interactions^17^. In addition, the TOC1-PRR domain was shown to have transcriptional repression potential in a Gal4-LexA/UAS system^5^. In *Arabidopsis*, PRR5, PRR7, and PRR9 function as transcriptional repressors by interacting with the plant Groucho/TUP1 co-repressor TOPLESS (TPL) through an EAR (ethylene-responsive element binding factor-associated amphiphilic repression) motif^18^. TPL recruits the histone deacetylase HDA19 and HDA6 to this protein complex and facilitates gene silencing^18^. However, TOC1 lacks this EAR motif and cannot interact directly with TPL. These observations suggest that TOC1 may function as a transcriptional repressor via a novel mechanism.

Previous studies have shown that full length TOC1 binds to a DNA motif (TGTG) in the promoter of *CCA1* directly through its CCT domain^5^. This TGTG motif is part of the previously defined GTGTGG “morning element” (ME) and CATGTG “hormone up-regulated at dawn motif” (HUD), which are found in the promoters of many morning genes. The CCT domain of CONSTANS (CO), a transcriptional activator for flowering^19,20^, was recently shown to bind a DNA motif (CORE2: GATTGTGGTTATGATTT) in the promoter of *FLOWERING LOCUS T (FT)* by interacting with two Nuclear Factor-Y transcription factors^21–23^. In addition, three elements were found to be enriched in the promoter of TOC1 target genes: G box (CACGTG), TBS (TCP binding site GGNCCCAC), and the plant-specific GA motif (AGARRGARRRAGADR)^5,13^. Consistently, it was found that TOC1 interacts with the phytochrome-interacting factor (PIF3 and PIF4)^24–26^ and CCA1 Hiking Expedition (CHE)^27^, which binds to the G box and TBS, respectively. Taken together, these results suggest that TOC1 can either bind to a DNA motif directly, or be recruited to the promoters of its targets by interacting with another transcription factor.

Although TOC1 has been identified as a transcriptional repressor, the biochemical properties of this protein are still unclear. In this study, we show that the CCT domain of TOC1 (TOC1-CCT) binds directly to RNA. We applied in vitro RNA selection to determine a G-rich RNA motif that is recognized by TOC1-CCT domain. Using quantitative electrophoretic mobility shift assays (EMSAs) and isothermal titration calorimetry (ITC), we showed that the TOC1-CCT domain binds RNA with nanomolar affinity. NMR spectroscopy showed that the RNA substrate interacts with lysyl and arginyl residues of the CCT domain. Mutations of these residues in the CCT domain resulted in either reduction or complete loss of RNA binding activity in vitro. Furthermore, the results in protoplasts showed that the residues involved in RNA binding are required for TOC1 to function as a transcriptional repressor in vivo. Our data strongly suggest that TOC1 activity might not be primarily as a canonical transcription factor, but instead play key roles as a multi-function transcriptional regulator.

## Results

### Identifying the TOC1-CCT consensus RNA target sequence

Previous studies have shown that the TOC1-CCT domain directly binds to double-stranded DNA (dsDNA) in vitro. However, the motifs enriched in TOC1 targets share low sequence similarity. To determine whether TOC1 is a site-specific DNA binding protein, we performed Systematic Evolution of Ligands by Exponential Enrichment (SELEX)^28^ with the purified TOC1-CCT domain (Supplementary Fig. 1) and random dsDNA. Interestingly, there was no significant enrichment of dsDNA motifs shown in SELEX (data not shown), suggesting that the TOC1-CCT domain binds to dsDNA with low affinity. The overexpression of TOC1 resulted in the disruption of expression of ∼2500 target genes^5^, but only a limited number of predicted targets were found to be occupied by TOC1^13^. Thus, we hypothesize that TOC1 may regulate its targets’ RNA abundance via additional mechanisms, e.g. by interacting with RNA directly. To investigate whether TOC1-CCT binds to RNA, we performed SELEX with the purified TOC1-CCT domain and random RNA (Fig. 1a). The purified TOC1-CCT domain was incubated with a pool of random undecamer RNA molecules (Fig. 1a). The eluted RNA fragments were pooled, amplified by reverse transcription PCR and transcribed for the next round of selection. After four rounds of selection, the RNA samples from the second and fourth rounds were collected and sent for deep-sequencing. A G-rich RNA motif (GGAGGGG) was significantly enriched in both samples (Fig. 1b).

**Fig. 1.**
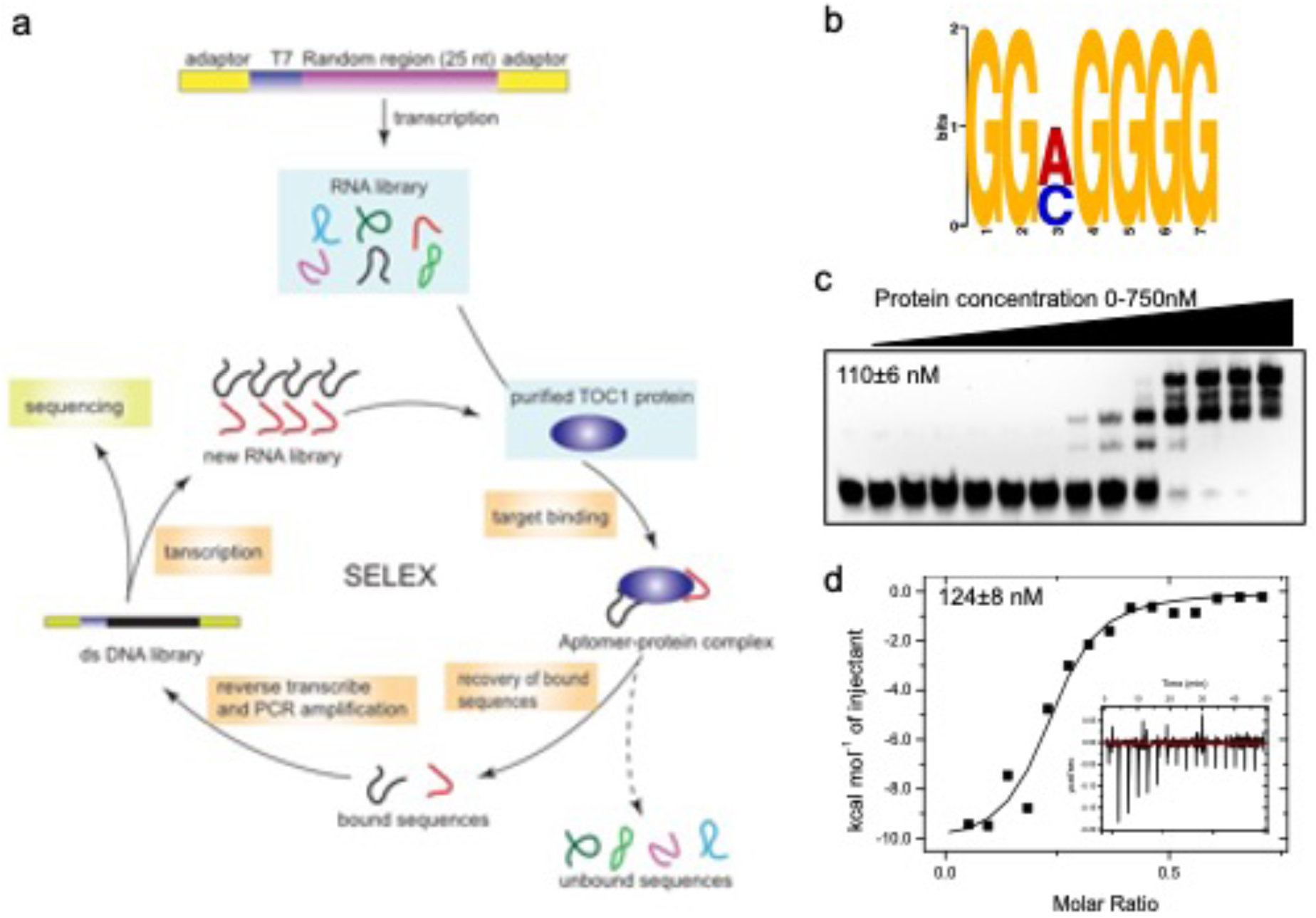
Identification of TOC1-CCT domain RNA target *in vitro*. (**a**) Schematic representation of SELEX procedure. (**b**) Weblogos of the most highly enriched motifs identified from round 2 and 4 of the TOC1-CCT SELEX dataset. (**c**) EMSA to assess the binding of TOC1-CCT domain to a biotin labeled G-rich RNA motif. Quantification of the binding affinity obtained a dissociation constant (*K*D) value of 110 ± 6 nM. (**d**) Isothermal titration calorimetry detection of interactions between TOC1-CCT domain and a G-rich RNA motif (these proteins tended to aggregate when added in excess of available RNA binding sites, giving rise to background signals at the end of the run) Quantification of the binding affinity obtained a dissociation constant (*K*D) value of 124 ± 8 nM determined by isothermal titration calorimetry (ITC).

To test whether the TOC1-CCT domain directly binds to this G-rich motif, we performed an electrophoretic mobility shift assay (EMSA) with biotin labelled RNA probes and purified TOC1-CCT domain (Fig. 1c). As shown in Fig. 1c, increasing concentrations of the TOC1-CCT domain resulted in the appearance of up-shifted RNA bands, indicating formation of an RNA-protein complex. The observed multi-bands with different sizes at higher concentrations of protein may be caused by the formation of TOC-CCT dimers or trimers with RNA probes. Quantification of the binding affinity obtained a dissociation constant (*K*D) value of 110 ± 6 nM determined by the EMSA and 124 ± 8 nM determined by isothermal titration calorimetry (ITC) (Fig. 1c and 1d).

### TOC1-CCT domain binds specifically to RNA

To determine whether the TOC1-CCT domain specifically binds to RNA, we performed a competition EMSA with unlabeled RNA. We observed that the TOC1-CCT domain binds to the G-rich RNA motif with different flanking sequences, and unlabeled RNA can outcompete biotin labeled G-rich RNA probes, suggesting that the RNA binding of TOC1-CCT domain is specific (Supplementary Fig. 2). To exclude the possible effect of the glutathione S-transferase (GST) tag, we performed an EMSA with the purified TOC1-CCT domain tagged with Phage late control gene D protein (GpD), GST-LUX ARRHYTHMO (LUX) and GST alone (Supplementary Fig. 3). These results clearly showed that only the TOC1-CCT domain binds to the G-rich RNA probe, not the LUX-GST or GST protein. To test whether the TOC1-CCT domain binds to other types of oligonucleotides containing the G-rich motif, we performed EMSA with RNA, DNA, double-stranded RNA (dsRNA), reverse RNA (rRNA) and a hybrid of RNA and ssDNA (Supplementary Fig. 4). Based on the intensity of the shifted bands we observed from all of the combinations, TOC1-CCT formed a less intense shifted band with DNA compared to the shifted band with RNA, indicating that the TOC1-CCT domain also has DNA binding activity. To assess the binding affinity, we determined apparent dissociation constants using EMSA with DNA and RNA in the presence or absence of various concentrations of TOC1-CCT domain (Supplementary Fig. 5). These results showed that the TOC1-CCT domain has a higher binding affinity for G-rich RNA.

**Fig. 2.**
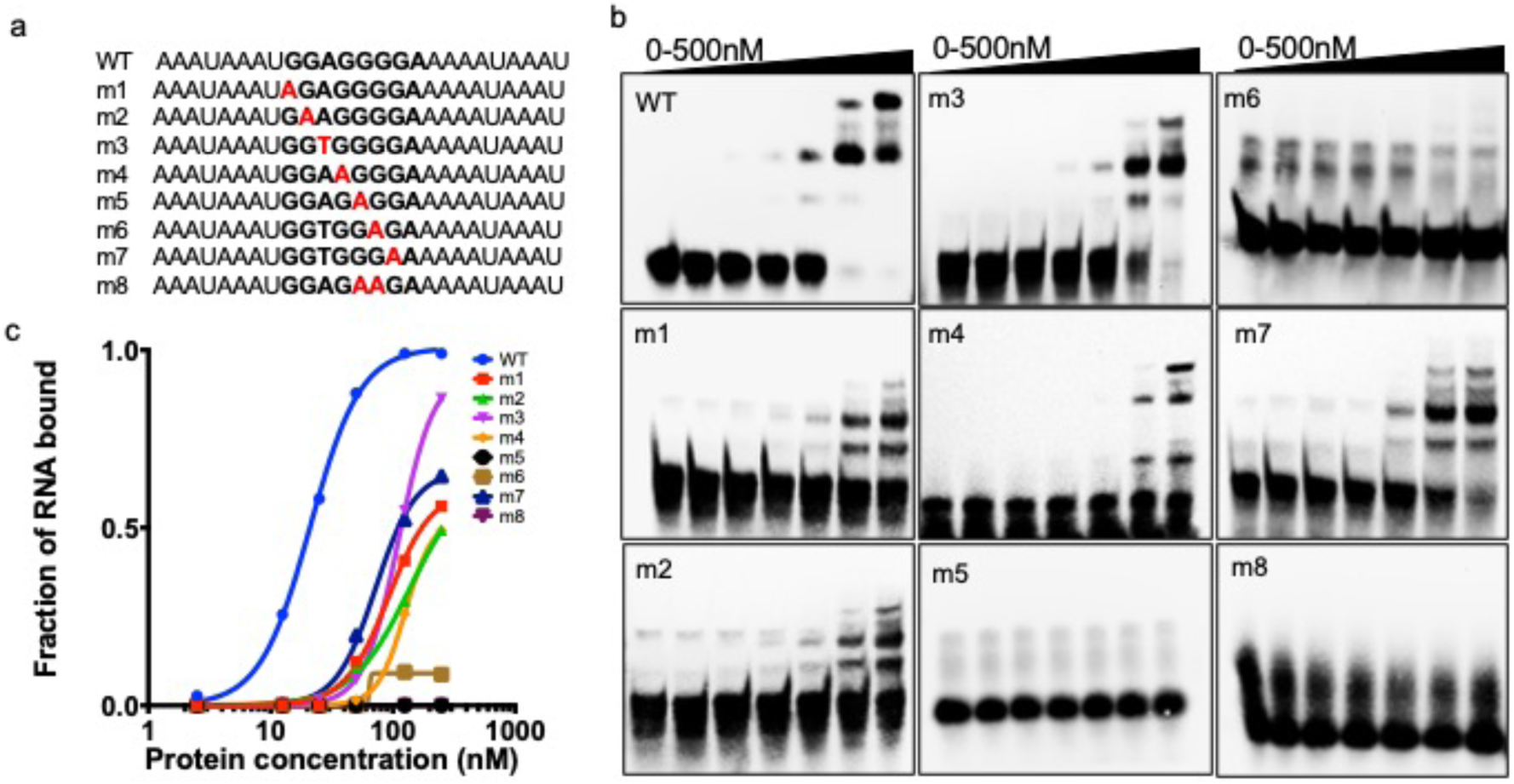
TOC1-CCT domain binds to a G-rich RNA motif in a sequence specific manner. (**a**) Nucleotide sequences of either wildtype or mutated RNA motif used for EMSA. The mutated nucleotides are highlighted. (**b**) Representative EMSAs of TOC1-CCT domain in the presence of probes with altered guanine to adenine or adenine to thymine in the G-rich motif. (**c**) Binding curves for the TOC1-CCT domain to RNA motifs.

**Fig. 3.**
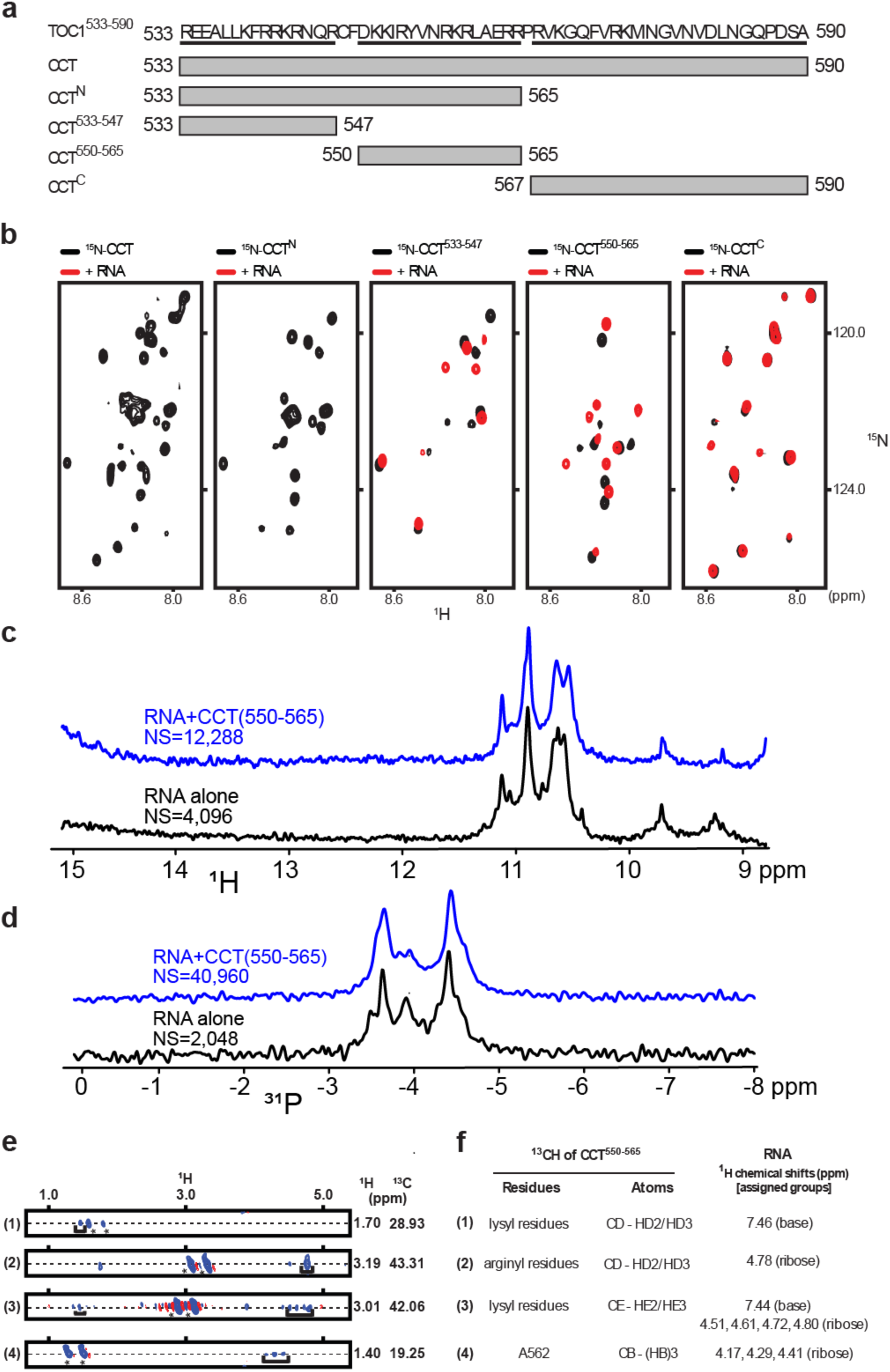
NMR characterization of TOC1-CCT-RNA interactions. (**a**)Schematic diagram of CCT constructs tested for binding RNA. (**b**)Zoomed-in regions of ^15^N,^1^H-HSQC spectra of ^15^N-labeled CCT fragments (CCT, CCT^N^, CCT^533-547^, CCT^550-565^, and CCT^C^) (black) and their respective mixtures with RNA (red). The zoomed-in region was based on the chemical shifts range of ^15^N-labeled CCT and its fragments. See Supplementary Fig. 9 for full spectra. All NMR samples had identical protein concentrations. Spectra were collected using identical acquisition parameters and plotted at the same contour level. Thus, absence of red peaks in panels 1 and 2 are due to CCT line broadening in the presence of RNA. (**c**) ^1^H NMR spectra of RNA ± CCT^550-565^ from 9-15 ppm. NS=number of scans for a given spectrum. (**d**) ^31^P NMR spectra of RNA ± CCT^550-565^. (**e**) Strip plots showing intermolecular NOE contacts between ^13^C-labeled CCT^550-565^ and unlabeled RNA (5’ UGG AGG GGA 3’). Asterisks indicate diagonal peaks from ^13^CH groups of CCT^550-565^. (**f**) Assignments of intermolecular NOEs between lysyl and arginyl groups of CCT^550-565^ and RNA. The bracketed peaks at 1.46 ppm (1) and 1.44 ppm (3) in (**e**) are likely to be aliased, arising from chemical shifts at 7.46 ppm and 7.44 ppm. The sweep width for the indirect ^1^H dimension was 6 ppm. Atom names follow the IUPAC-IUBMB-IUPAB recommendations^57^.

**Fig. 4.**
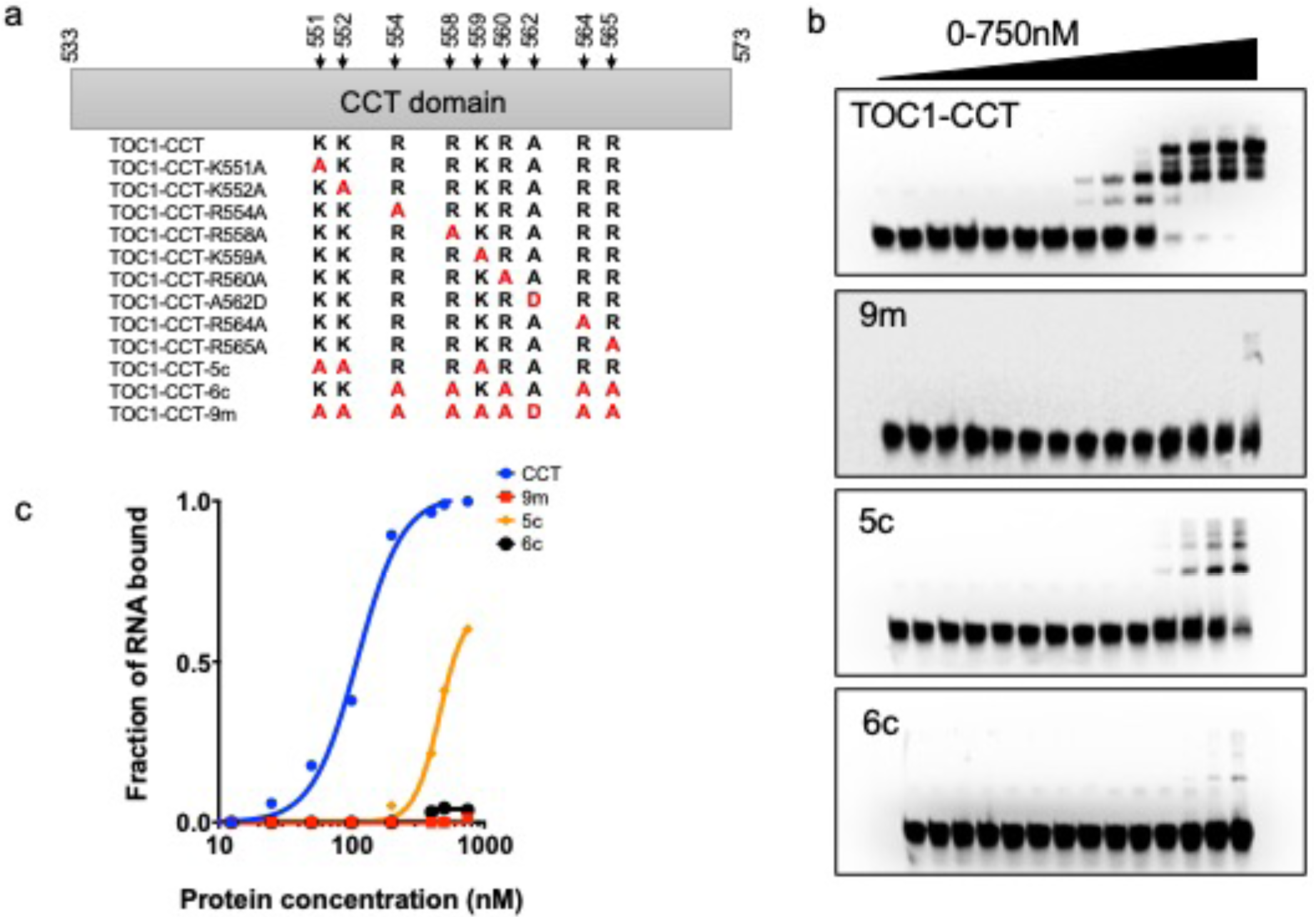
Mutational analysis of TOC1-CCT domain and RNA binding. (**a**) Diagram of the TOC1-CCT domain (amino acid 533-573). The identified residues from NMR are marked with black arrow. TOC1-CCT variants for EMSA in Fig. 4B are listed and mutated residues are marked with red color (**b**) EMSA experiments with the TOC1-CCT mutants 9m (K551A, K552A, R554A, R558A, K559A, R560A, A562D, R564A, R565A), 5c (K551A, K552A, K559A), 6c (R554A, R558A, R560A, R564A, R565A). (**c**) Binding curves of TOC1-CCT mutants from (**b**).

**Fig. 5.**
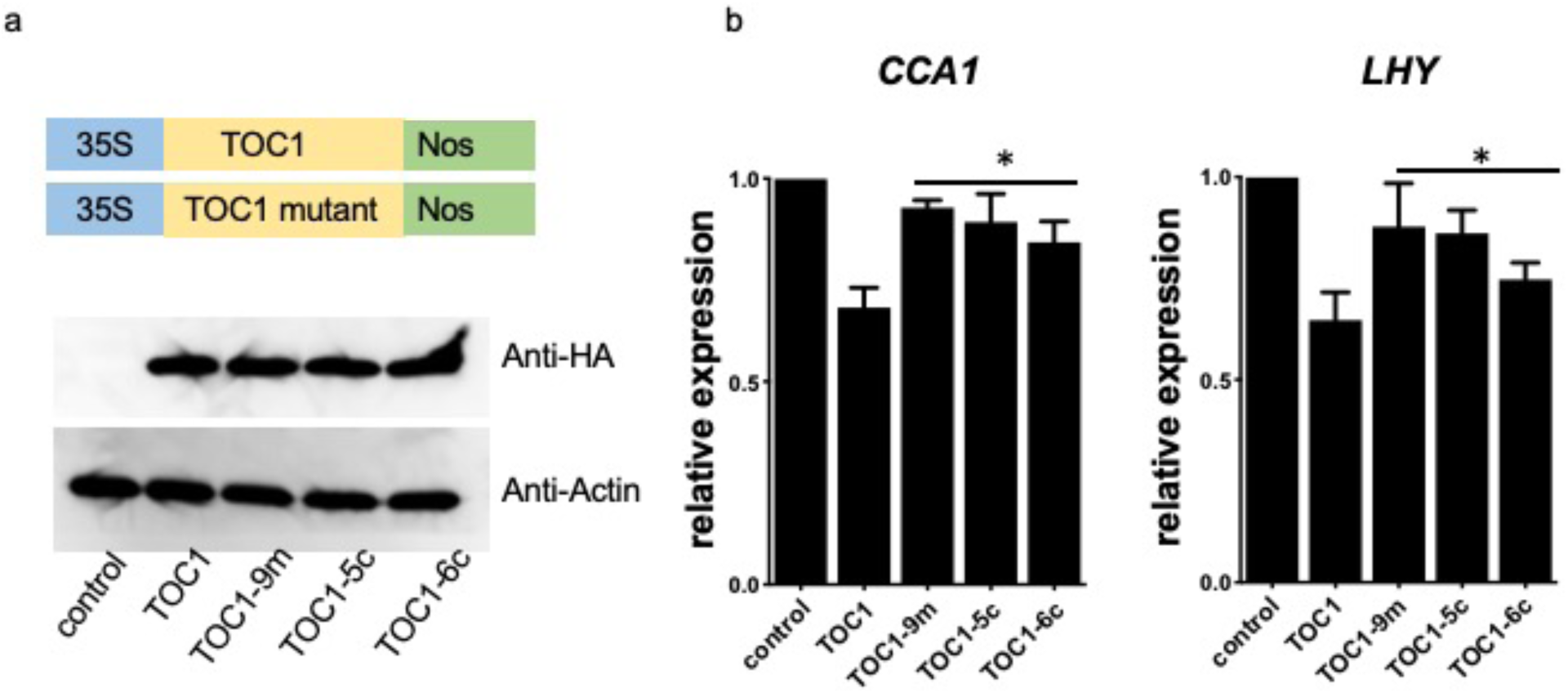
Functional analysis of TOC1-CCT domain mutations *in vivo*. (**a**) Immunoblot analysis of the expression levels of TOC1 and different TOC1 mutants, 9m (K551A, K552A, R554A, R558A, K559A, R560A, A562D, R564A, R565A), 5c (K551A, K552A, K559A), and 6c (R554A, R558A, R560A, R564A, R565A) in mesophyll protoplast. (**b**) Relative expression of *CCA1* and *LHY* in protoplast overexpressing TOC1 or different TOC1 mutants. Statistical significance was calculated by one-way ANOVA (*P < 0.05).

CCT is a conserved domain found in ∼40 proteins in the *Arabidopsis* genome. Based on the results of the TOC1-CCT domain, it is reasonable to speculate that the CCT domain has RNA-binding properties. Therefore, to investigate whether other CCT domains also exhibit similar RNA binding activity, we incubated the CCT domain from PRR5 and CO with biotin labeled G-rich RNA probes (Supplementary Fig. 6). The EMSA results showed that all three CCT domains (TOC-CCT, PRR5-CCT, CO-CCT) bind to the G-rich RNA probes, but with weaker binding affinities.

To further understand the importance of sequence context within RNA, we investigated the contribution of individual nucleotides in the G-rich RNA element for complex formation. We performed EMSA experiments with a number of RNA probes that contained individual substitutions of guanine to adenine or adenine to thymine in the G-rich motif (Fig. 2a). In comparison with the obvious shifted band seen using the wildtype probe, complex formation was almost abolished upon substitution of the guanine to adenine. In addition, changing the 3^rd^ base from adenine to thymine resulted in only a moderate reduction of protein-RNA interaction (Fig. 2b and 2c), demonstrating that the guanine is essential for the binding of TOC1-CCT domain to RNA.

### NMR shows that basic residues of CCT interact with RNA

The lack of precedence for any clock gene directly binding to RNA motivated us to examine other methods to validate interactions suggested by EMSA. As NMR is a sensitive tool in probing intermolecular interactions, we sought to conduct an NMR study of CCT-RNA interactions. Various CCT fragments (Fig. 3a) were designed based on PSIPRED (http://bioinf.cs.ucl.ac.uk/psipred/) secondary structure prediction (Supplementary Fig. 8), which suggested that CCT^533-547^ and CCT^550-565^ have *a*-helical propensity while a C-terminal fragment, CCT^C^ (CCT-567-590), has *b*-strand propensity. Under optimized binding conditions (10 mM Tris, 50 mM KCl, 5 mM MgCl2, pH 7.0), ^15^N,^1^H-HSQC^29^ (heteronuclear single quantum coherence) peaks of ^15^N-CCT were significantly broadened upon addition of RNA (5’ UGG AGG GGA 3’), as shown in Fig. 3b. The N-terminal fragment CCT^N^ (CCT-533-565) experienced similar NMR line broadening. Two sub-fragments of CCT^N^, CCT^533-547^ and CCT^550-565^, showed broadening to a lesser extent. ^15^N-labeled SUMO (data not shown) and CCT^C^ showed only weak chemical shift perturbation by RNA. Thus, the N-terminal half of CCT forms most of the interactions with RNA. We observed proton resonances between 10.5-11.5 ppm (Fig. 3a) in RNA motif, which are characteristic of H1 imino protons of guanosyl residues of G-quadruplexes. The presence of CCT^550-565^ significantly broadened the guanosyl H1 imino resonances (Fig. 3c) and ^31^P resonances of the RNA motif (Fig. 3d). Gel filtration analysis indicated that CCT-RNA complexes do not form aggregates (Supplementary Fig. 7).

Using a three-dimensional ^13^C-edited NOESY experiment^30,31^, we obtained intermolecular NOEs (Fig. 3e) between ^13^C-labeled CCT^550-565^ and unlabeled RNA. Backbone and sidechain assignments were achieved based on spectra from 3D triple resonance NMR experiments, including HNCACB^32^, HNCOCACB^33^, HNCO^34^, and HNCACO^35^, CCCONH^36^, HCCCONH^36^. The majority of NOE contacts were assigned to arginyl and lysyl residues of CCT^550-565^ (Fig. 3e), which probably interact with the negatively charged phosphate backbone of RNA. Interestingly, one contact originates from alanine 562 (A562) of the CCT, whose mutation to valine caused shortened circadian rhythms in light-grown *Arabidopsis*^4,37^. To model the CCT structure when bound to RNA, ^15^N,^13^C-labeled CCT chemical shifts were used as restraints for CS-Rosetta^38^ to generate structural models. The calculated CS-Rosetta models of CCT^550-565^ in this complex are largely unstructured (Supplemenatary Fig. 10).

### Mutational analysis of the protein-RNA interaction

The NMR experiments identified nine residues likely to be involved in the interaction between the TOC1-CCT domain and RNA. To examine the role of these residues in RNA binding, we substituted these residues individually for alanyl or aspartyl residues and tested RNA binding activity of the resulting mutants (Fig. 4a). Among these single amino acid mutants, K552A, R560A and A562D showed mildly impaired RNA binding activity (Supplementary Fig. 11). However, none of the single amino acyl mutants showed significant change in RNA binding activity as shown in Supplementary Fig. 11. This result suggested that multiple residues from TOC1-CCT may be involved in the complex formation with RNA. To test this hypothesis, we used site-directed mutagenesis to construct a TOC1-CCT-9m mutant (Fig. 4a), in which all nine residues were mutated to alanine or aspartic acid. To test whether TOC1-CCT-9m retains RNA binding activity, we performed EMSA experiments with a biotin-labelled RNA probe. As shown in Fig. 4b and 4c, TOC1-CCT-9m lost almost all RNA binding activity, indicating that these residues bind to RNA in a cooperative manner. We further divided the nine residues into two groups according to their biochemical properties: TOC1-CCT-6c (R554A-R558A-R560A-R564A-R565A) and TOC1-CCT-5c (K551A-K552A-K559A). As shown in Fig. 4b and 4c, both TOC1-CCT-6c and TOC1-CCT-5c also displayed a significant reduction in RNA binding activity.

### RNA binding properties of TOC1-CCT is required for target regulation in vivo

Given that the identified residues are critical for in vitro RNA binding, we further tested the importance of these residues for TOC1 function in vivo. We expressed wild type TOC1 protein and mutants containing single or multiple mutations in mesophyll protoplast (Fig. 5a). Using qRT-PCR, we found that wildtype TOC1 effectively down-regulated *CCA1* and *LHY* expression. Single amino acid changes in TOC1 resulted in only mild reduction on *CCA1* and *LHY* repression (Supplementary Fig. 12), consistent with their effects on RNA binding activity in vitro. This result suggested that these RNA binding residues may work cooperatively. To test this possibility, we used sited-directed mutagenesis to construct a full length TOC1-9m mutant, in which all nine residues were mutated. In comparison with the single mutants (Supplementary Fig. 12), TOC1-9m protein caused an obvious reduction in *CCA1* and *LHY* repression (Fig. 5b), consistent with the in vitro observation that TOC1-CCT-9m mutant caused significantly reduced affinity with G-rich RNA. Similarly, overexpressing TOC1-5c and TOC1-6c in protoplasts resulted in a reduction of *CCA1* and *LHY* repression (Fig. 5b). Based on these findings, we conclude that the RNA binding activity of TOC1 in vitro is directly related to its role in target regulation in vivo.

## Discussion

As a master regulator of many aspects of plant life, clock proteins need to incorporate properties of transcription, post-transcription and translation to fine tune the regulation of their targets throughout the day. A role for clock proteins’ involvement with RNA indirectly has been reported previously. In *Arabidopsis*, the clock output gene *Glycine Rich Protein 7 (GRP7)* has features of an RNA binding protein and has been shown to be involved in mRNA splicing^39^. Outside of plants, the *Neurospora* circadian clock gene *Frequency (FRQ)* associates with FRQ-interacting RNA helicase (FRH), an essential DEAD box containing RNA helicase that is important for the stability and complex formation of FRQ and its regulatory role in the *Neurospora* clock^40^. However, its direct function in mRNA processing requires further clarification. In none of these studies has direct association of a clock protein with RNA been reported. Using a combination of biochemical studies, reporter assays, and structural analyses by NMR, we provide evidence supporting a role for TOC1 as a RNA binding protein.

Since its discovery, dissecting the biochemical and transcriptional regulatory properties of TOC1 in the clock has proven to be challenging. Overwhelming and opposing experimental evidence for TOC1 transcriptional regulation (binding motif and targets) suggested that the biological function of this founding and core clock protein might be complex. Using SELEX, we revealed that the TOC1-CCT domain directly binds to a G-rich RNA motif. Both EMSA experiments and an ITC assay indicated high binding affinity of the TOC1-CCT domain with RNA. Unlabeled RNA competitors compete biotin-labeled G-rich RNA probes from the TOC1-CCT domain, suggesting the RNA binding of the CCT domain is specific. Additionally, the CCT domain from CO and PRR5 also exhibit RNA binding activity, indicating that the conserved CCT domain and RNA binding properties are functionally relevant. Furthermore, the CCT domain also displayed lower binding affinity to DNA, indicating that this domain may bind DNA and RNA in vivo. In fact, recent studies have revealed that several proteins also exhibit such dual binding activity. For example, in mammalian systems, transactive response (TAR) DNA-binding protein 43 (TDP-43) has been shown to bind thousands of RNA in neurons^41^ and a RNA binding protein Lin28A also binds to DNA directly in mice^42^.

RNA mutations that had the most severe effects on TOC1-CCT’s RNA binding activity (more than 100-fold reduction in affinity) resulted from guanine-to-adenine substitutions. Mutating adenine to thymine in the RNA motif had little effect on TOC1-CCT domain binding, suggesting that guanine is essential for the RNA binding activity of the CCT domain. Interestingly, G-rich motifs were significantly enriched from CLIP-seq data (crosslinking immunoprecipitation followed by deep sequencing) for several mammalian RNA binding proteins^43,44^. These RNA binding proteins regulate the abundance of their target RNAs by controlling RNA processing, such as alterative splicing and polyadenylation^43^. Thus, identifying the RNA targets of TOC1 by genome-wide methods such as RNA Immunoprecipitation followed by sequencing (RIP-seq) will help us to understand how TOC1 regulates its targets via an RNA mechanism.

NMR experiments showed binding between the CCT and G-rich RNA, consistent with the EMSA experiments. In addition, the NMR experiments revealed two minimal fragments CCT^533-547^ and CCT^550-565^ required for binding RNA. The NOE study with CCT^550-565^ identified up to nine residues making contact with RNA. The CS-Rosetta program suggested that CCT^550-565^ in the bound state is largely unstructured. The many basic residues in the N-terminal half of CCT suggest that CCT makes several electrostatic contacts with the acidic backbone of RNA. EMSA experiments showed that CCT binding is sensitive to nucleotide sequence, whereas NMR data suggest electrostatic contacts with the monotonous RNA backbone. Taken together, it is likely that full-length CCT specifically recognizes G-quadruplex structures through charge-charge interactions.

Intermolecular contacts between CCT^500-565^ and RNA identified by NMR guided mutagenesis studies on CCT-RNA binding and functional validation in vivo. We observed that single amino acyl substitutions in the TOC1-CCT domain showed only minimal reduction in RNA binding (Supplementary Fig. 11). Interestingly, the mutation at A562, originally identified in the *toc1-1* mutant, also reduced RNA binding here (Fig. 3c and Supplementary Fig. 11). Moreover, selected combinations of mutations of the nine RNA-interacting residues of CCT resulted in either a severe or complete loss of RNA binding activity in vitro, implying that the TOC1-CCT domain binds to RNA in a cooperative manner (Fig. 4a). Over-expression of full length TOC1 in protoplasts resulted in down-regulation of *CCA1* and *LHY* expression, while TOC1 mutants had a reduced effect, suggesting that RNA binding of the CCT domain is required for TOC1 function as a transcriptional repressor in vivo. In addition, TOC1 has been shown to interact with four RNA binding proteins (RBPs) in an *Arabidopsis* interactome study^45^. These RBPs are rhythmically expressed and may form a putative ribonucleoprotein (RNP) complex with TOC1 to regulate its target RNA processing and metabolism. Thus, comprehensive identification and characterization of the RNA binding proteins that interact with TOC1 will help us to understand the regulatory role of this founding clock protein at the RNA level.

In summary, our work provides the first scenario in which TOC1 could bind to RNA and establishes a molecular connection with its regulatory function. Our findings provide a novel role for TOC1 that paves the way to further investigate alternative mechanisms underlying circadian clock regulation of gene expression.

## Material and Methods

### Expression and purification of recombinant proteins

The TOC1-CCT domain (residues 533-575), PRR5-CCT domain (residues 509-551) and CO-CCT domain (residues 306-348) were amplified from the *Arabidopsis* TF library^46^ and cloned into a bacterial expression plasmid based on the backbone from pET28b and pET42b vectors. The GpD tag was added at the N terminal of the pET28b vector. Mutations were introduced into the CCT domain by site-directed mutagenesis (Pfu Turbo DNA polymerase, Agilent) and all the constructs were verified by sequencing and subsequently transformed into BL21-CodonPlus (DE3)-RIL Competent Cells for protein production.

1ml of bacterial culture was grown overnight at 30°C and inoculated into 500ml TB medium with ampicillin (100 μg/mL) and chloramphenicol (30 μg/mL) at 37°C for 3-4 hours to an OD600 of 0.4 to 0.6. Protein expression was induced with 0.4 mM isopropyl β-D-1-thiogalactopyranoside (IPTG) for 5hr at 30°C. Bacterial cells (∼10g) were collected (3000 rpm, 20 min, 4°C) and lysed in Buffer A (50 mM Tris pH=8.0, 1 M NaCl, 1 mM DTT, 1 tablet of protease inhibitor, 20 mM imidazole and 1 mg/ml lysozyme) for 30min in the cold room (4°C). The soluble fraction of lysates was incubated with TALON resin (clontech) for 30min and eluted with 200ul Buffer B five times (50 mM Tris pH=8.0, 1 M NaCl, 1 mM DTT, 1 tablet of protease inhibitor and 250 mM imidazole). The purified proteins were exchanged into PBS buffer with 10% glycerol using PD SpinTrap G-25 columns (GE), and concentrated with protein concentrators (Vivaspin 500, MWCO 3K, GE). The protein samples for ITC were dialyzed overnight with 4L PBS buffer in the cold room.

### Systematic evolution of ligands by exponential enrichment experiments (SELEX)

SELEX was performed using TOC1-CCT proteins. Initially, a random library of oligonucleotides was converted into double strand DNA and transcribed into RNA as described (James L. Manley 2011). For each selection cycle, 1μM protein was mixed with 10nM RNA in SELEX buffer (20 mM Tris-HCl pH=8.0, 250 mM NaCl, 2 mM MgCl_2_, 1 mM DTT and 1% glycerol) for 15 min. The RNA libraries obtained after four cycles of selection were subsequently used for high-throughput sequencing. The unique most frequent sequences were aligned using MEME software.

### In Vitro RNA-Binding Assays

Unlabeled and 5′ biotin-labeled RNA (AAAUAAAUGGAGGGGAAAAAUAAAU) were purchased as synthetic oligonucleotides from IDT (Integrated DNA Technologies). All RNA probes were purified using spin columns and precipitated by ethanol and re-suspended in RNase-free water prior to use. Biotin labeled RNAs were mixed with recombinant proteins in RNase-free buffer (20 mM Tris-HCl, pH=8.0, 100 mM NaCl, 4% glycerol, 1 mM DTT) in a final volume of 20μl for 30 min at 4°C. Then, the samples were separated in a 10% polyacrylamide gel buffered with 0.5 × TBE at 4 °C. Next, the separated proteins on the gel were transferred onto a positively charged nylon membrane (HybondTM-XL) at 380 mA for 40 min. The membrane was crosslinked in a UV crosslinker (254nm) at 120 mJ/cm2 for 45 seconds and incubated with stabilized streptavidin-HRP for 15 min. The membrane was washed three times and exposed to a CCD camera for 5 min. Apparent *K*D values were obtained by fitting the resulting data points into a single exponential Hill using GraphPad Prism (GraphPad Software, Inc., La Jolla, CA).

### NMR

CCT constructs were cloned into the pET-28b vector as *SUMO* fusion genes using NdeI and HindIII sites. The protein expression protocol for NMR samples including ^15^N- or ^15^N,^13^C-labeled CCT or its fragments has been described previously^47^, except for using the BL21-CodonPlus(DE3)-RIPL strain (Agilent) and supplementing additional chloramphenicol (50 µg/mL) for initial cultures. Cells were resuspended in *Lysis Buffer* [50 mM NaH_2_PO_4_, 500 mM NaCl, pH 8.0] and lysed using an EmulsiFlex-C55 homogenizer (Avestin). Lysates were clarified at 13,000 rpm for 30 min and the resulting supernatants were subjected to specific purification procedures described below.

^15^N-labeled SUMO fusion CCT or its fragments, which were tested for binding RNA (5’ UGG AGG GGA 3’) (GE Dharmacon), were purified by Nickel affinity chromatography [*Wash Buffer*: 50 mM NaH_2_PO_4_, 500 mM NaCl, 20 mM imidazole, pH 8.0; *Elution Buffer*: 50 mM NaH_2_PO_4_, 500 mM NaCl, 250 mM imidazole, pH 8.0; Elution volume: 5 mL] and gel filtration chromatography [*Elution Buffer*: 10 mM Tris, 50 mM KCl, pH 7.0]. Ulp1 stocks at 100 µM in the buffer [25 mM NaH_2_PO_4_, 250 mM NaCl, 125 mM imidazole, 50% glycerol, pH 8.0] were added (final concentration: 1 µM) into the SUMO fusion proteins (CCT or its fragments) for sufficient cleavage before collecting NMR spectra. The Ulp1 cleavage at room temperature took about 20 minutes for ^15^N-labeled SUMO fusion proteins of CCT^550-565^ and CCT^567-590^, and 5 hours for ^15^N-labeled SUMO fusion proteins of CCT, CCT^533-565^, and CCT^533-547^, respectively. As a control, Ulp1 was added into ^15^N-labeled SUMO and the sample was incubated at room temperature for about 20 minutes. See Supplementary Table S1 for NMR sample details. Note that protein concentrations were determined using the Bradford assay unless otherwise indicated.

Two D_2_O NMR samples [^15^N,^13^C-CCT^550-565^ + ^15^N,^13^C-SUMO + RNA in 99.8% D_2_O; ^15^N,^13^C-CCT^550-565^+ RNA] were prepared for obtaining intermolecular NOE contacts between CCT^550-565^ and RNA.

- CCT^550-565^ for the sample which contains SUMO impurities [^15^N,^13^C-CCT^550-565^ +^15^N,^13^C-SUMO + RNA; 99.8% D_2_O] was purified by Ni affinity chromatography as described above, followed by Ulp1 cleavage for >12 h at 4 °C [addition of 100 µL of Ulp1 stock (100 µM) into], separation from Ulp1 and SUMO by a 10-kDa cutoff disc using the Amicon concentrator, and buffer exchange into the NMR sample buffer [10 mM Tris, 50 mM KCl, 5 mM MgCl_2_, pH 7.0]. The protein sample was lyophilized and redissolved in 99.8% D_2_O. The protein concentration was estimated by using the Bradford assay. RNA stock in the buffer [10 mM Tris, 50 mM KCl, 5 mM MgCl_2_, pH 7.0, 99.8% D_2_O] was mixed with the protein sample and supplemented with DSS to a final concentration of 20 µM. The NMR sample looked very cloudy upon mixing but cleared up after incubation at room temperature for about 2 h. Clarification at 13,000 rpm for 2 min shows only a small pellet. The clarified sample was used for NMR data collection. The same incubation and clarification protocol was followed for subsequent sample preparations.
- CCT^550-565^ for the pure sample [^15^N,^13^C-CCT^550-565^ + RNA; 99.96% D_2_O] was purified by Ni affinity chromatography, Ulp1 cleavage, as described above, buffer exchange into the low salt buffer [Buffer A: 50 mM NaH_2_PO_4_, pH 7.0], cation exchange chromatography [*Column*: HiTrap SP HP 5 mL; *Buffer A*: 50 mM NaH_2_PO_4_, pH 7.0; *Buffer B*: 50 mM NaH_2_PO_4_, 1 M NaCl, pH 7.0], and buffer exchange into the NMR sample buffer [10 mM Tris, 50 mM KCl, 5 mM MgCl_2_, pH 7.0]. The purity for this sample was over 95%. The protein stock was determined by A280 to be ∼1.1 mM. Bradford assay was not used for this pure sample, because it may underestimate the concentrations of peptides smaller than 3 kDa^48^ (molecular weight of CCT^550-565^: 2.1 kDa). The NMR sample was made by mixing RNA, DSS, and NaN3 with the 1.1 mM protein stock, lyophilized and redissolved in 99.96% D_2_O.

For backbone and assignment, an H2O sample [^15^N,^13^C-CCT^550-565^ + RNA; 95% H_2_O/5% D_2_O] was made under the same condition for its D_2_O counterpart, except with an H_2_O buffer. NMR experiments ^15^N,^1^H-HSQC^29^, HNCACB^32^, HNCOCACB^33^, HNCO^49^, and HNCACO^50^ were carried out on the H2O sample for residue-specific backbone assignments (^1^H^N, 15^N, ^13^C*a*, ^13^C*b*, ^13^C’). NMR experiments ^13^C,^1^H-HSQC^51^, CCCONH^36^, HCCCONH^36^ were carried out on this sample for residue-specific assignment of side-chain chemical shifts.

All NMR experiments were performed at 25 °C on a Bruker 600 MHz AVANCE III spectrometer equipped with a TCI cryoprobe with z-axis pulsed-field gradient capability. Data analysis was performed by using NMRPipe^52^, PINE-Sparky^53^, and MARS^54^. Structural models of CCT^550-565^ in its complex with RNA were generated by CS-Rosetta^38^. See Supplementary Tables S2 and S3 for NMR parameters.

### Protoplast Isolation and transformation

For leaf mesophyll protoplast isolation, *Arabidopsis* Columbia WT plants were grown in a 12 h light / 12 h dark diurnal cycle with 70 μE light intensity for 4 weeks. Protoplast isolation was performed as described^55^. The protoplasts were transformed with a plasmid encoding wild-type TOC1 or its derivatives. The transformed cells were re-suspended in a WI solution containing 5% fetal bovine serum. After six hours’ incubation, the protoplasts were harvested (100rpm, 3min) at room temperature.

### qRT-PCR and Western blotting analyses

Total RNA was isolated with the Qiagen RNeasy plant mini kit (Qiagen). cDNA was synthesized using 1 µg of total RNA and reverse-transcribed with the iScript cDNA synthesis kit (Bio-Rad). Synthesized cDNAs were then quantified by real time quantitative PCR (qPCR) as described previously^56^. The primers used to quantify the expression of *LHY* were 5’ ACCAACGAAACAGGTAAGTGGCG 3’ and 5’ GTGCACAGTAGTACCATCGTTACC 3’, and for *CCA1* were 5’ CCGCAACTTTCGCCTCAT 3’ and 5’ GCCAGATTCGGAGGTGAGTTC 3’. As a normalization control, we used *isopentenyl pyrophosphate:dimethylallyl pyrophosphate isomerase* (*IPP2*). PCR conditions were as follows: 95°C for 3 minutes followed by 40 cycles of 95°C for 10 seconds, 55°C for 15 seconds and 72°C for 15 seconds.

For protein analysis, protoplast samples were lysed with 40 μl loading buffer (Tris-HCl 50 mM, pH=8.2, glycerol 20%, protease inhibitors), boiled for 5 min at 95°C, then loaded on a 10% SDS-PAGE gel and separated in 1x MOPS running buffer at 120 Volts for 1 h. After separation, proteins were transferred to a PVDF membrane (Immobilon®-P, Millipore) with a semi-dry transfer system (Trans-Blot® SD, Bio-Rad) in 1x MOPS buffer with 10% methanol at 12 Volts for 1 h. After blocking with 5% skim milk, the membrane was incubated with antibody in 1% milk for 2 hours. Then, the membrane was washed 5 times in TBST (50 mM Tris, 150 mM NaCl, 0,05% Tween 20), incubated with Pierce SuperSignal® West Pico chemiluminescent substrate (Thermo Scientific, 34078) for 1 min and exposed to film for 1 min.

### Isothermal titration calorimetry (ITC)

The purified proteins were extensively dialyzed at 4°C in 4L PBS buffer using dialysis tubing with molecular cutoff at 3.5 kDa (SnakeSkin Dialysis Tubing, 3.5K MWCO, 22 mm, Fisher). ITC measurements were performed on a MicroCal ITC200 (GE Healthcare) at 25°C. Protein/RNA concentration for the cell and syringe for the ITC assays were 50 μM protein into 50 μM RNA. All data were best fit using Origin 7.

## Acknowledgements

We thank Susanna Wang for critical reading of the manuscript, Ian Wilson for supporting ITC, and members of the Kay and LiWang labs for helpful discussions. Research reported in this publication was supported by the National Institutes of General Medicine of the National Institutes of Health under award numbers R01GM056006, R01GM067837 to S.A.K, and R01GM107521 to A.L. We thank Steve Grimaldi for maintaining the UCM NMR cryoplatform and David Rice maintaining the UCM NMR Facility.

The content is solely the responsibility of the authors and does not necessarily represent the official views of the National Institutes of Health.

## Authors Contribution

LY, YGC, DHN, AL and SAK designed the experiments, LY and YGC performed the experiments, LY, YGC, DHN, CHG, AL and SAK analyzed the data, and LY, YGC, DHN, AL and SAK wrote the manuscript.

The authors declare no conflict of interest.

## Notes

### Competing Interest Statement

The authors have declared no competing interest.

