## supplementary Figures for "A Novel Regulatory Role for the Circadian Clock Protein TOC1 via RNA binding"

**Supplementary Fig. 1** TOC1-CCT domain purification.

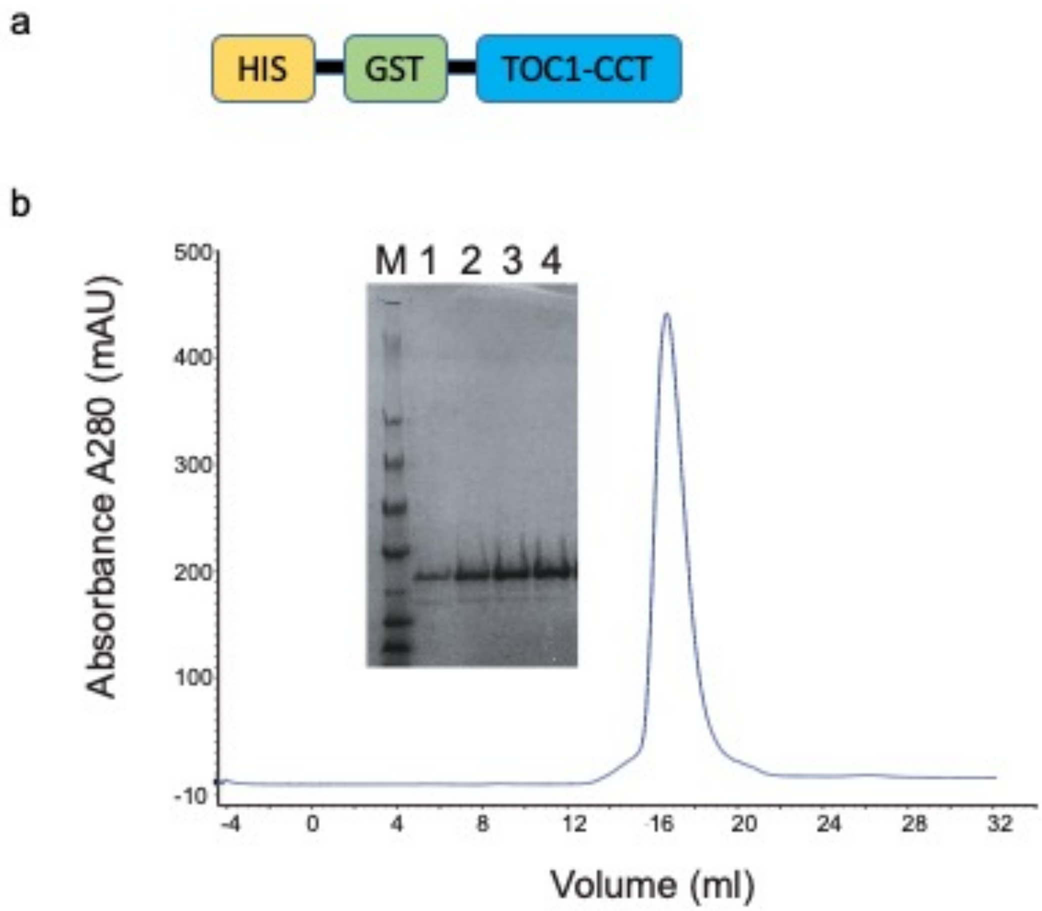


**Supplementary Fig. 1**

TOC1-CCT domain purification. (**a**) Schematic of recombinant TOC1-CCT domain tagged with GST and His. (**b**) Coomassie-stained gel of purified TOC1-CCT domain and analysis by size exclusion column.

**Supplementary Fig. 2** TOC1-CCT binds specifically to a G-rich RNA motif.

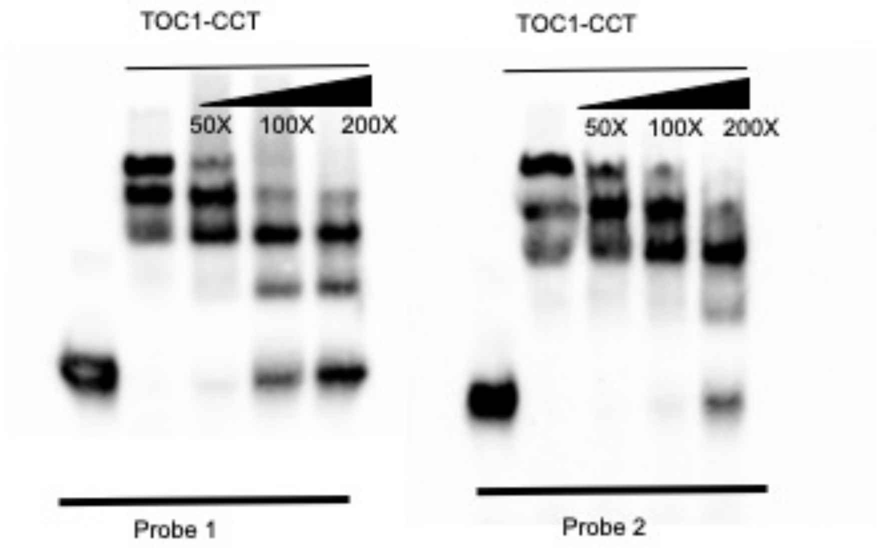


**Supplementary Fig. 2**

TOC1-CCT binds specifically to a G-rich RNA motif. EMSA showing recombinant TOC1-CCT binds to two RNA probes containing a G-rich motif. Respective unlabeled probes with the same sequence as the biotin-labeled probes were used as competitors.

.

**Supplementary Fig. 3** TOC1-CCT, but not GST and GST-LUX, binds to RNA

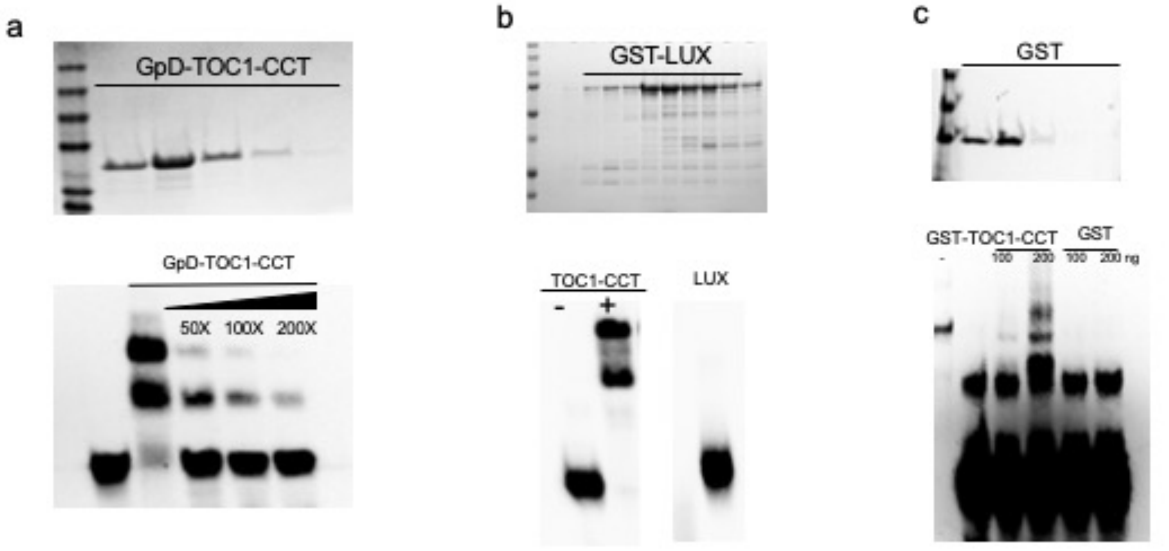


**Supplementary Fig. 3**

TOC1-CCT, but not GST and GST-LUX, binds to RNA. (**a**) Purification of TOC1-CCT domain tagged with GpD domain. EMSA of the binding of the recombinant TOC1-CCT-GpD to RNA probes containing a G-rich motif. (**b**) and (**c**) Purification of LUX-GST and GST protein. EMSA of the binding of the recombinant purified protein to RNA probes containing G-rich motif.

**Supplementary Fig. 4** EMSA showing recombinant TOC1-CCT binds to G-rich DNA and RNA.

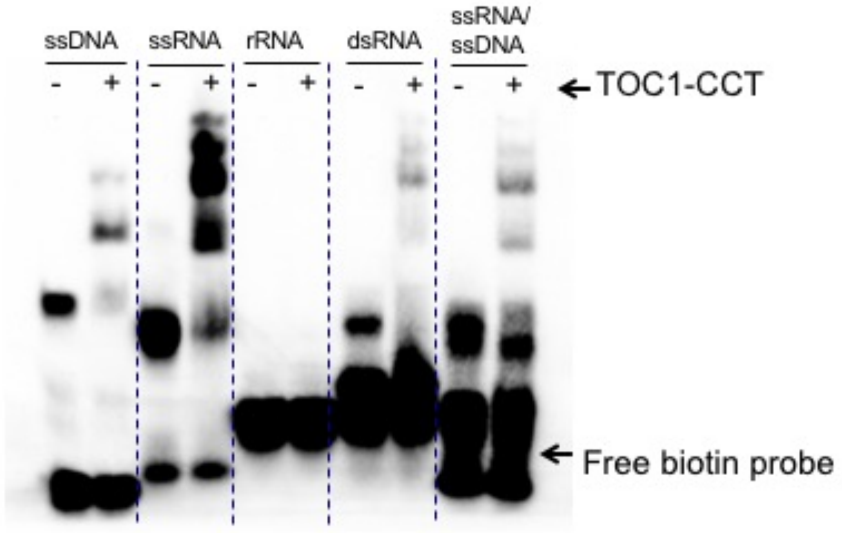


**Supplementary Fig. 4**

EMSA showing recombinant TOC1-CCT binds to G-rich DNA and RNA.

**Supplementary Fig. 5** TOC1-CCT binds to DNA and RNA.

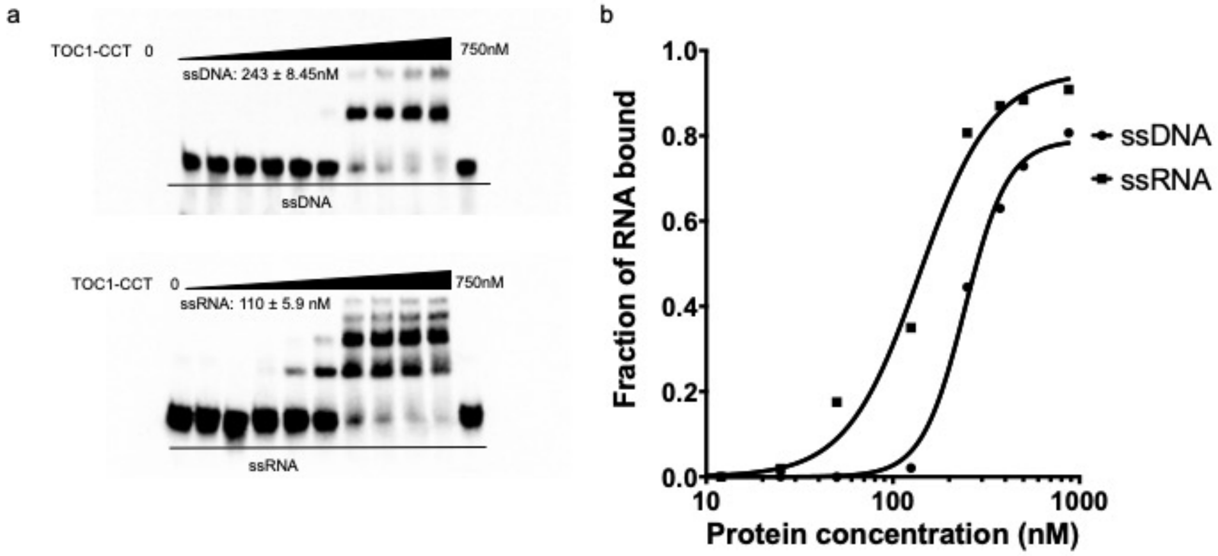


**Supplementary Fig. 5**

TOC1-CCT binds to DNA and RNA. (**a**) EMSA showing recombinant TOC1-CCT binds to ssDNA and RNA. (**b**) Complete binding curves for TOC1-CCT domain to a G-rich RNA motif. The data were fitted by using a single exponential Hill equation.

**Supplementary Fig. 6** GST-tagged TOC1/CO/PRR5-CCT binds to RNA.

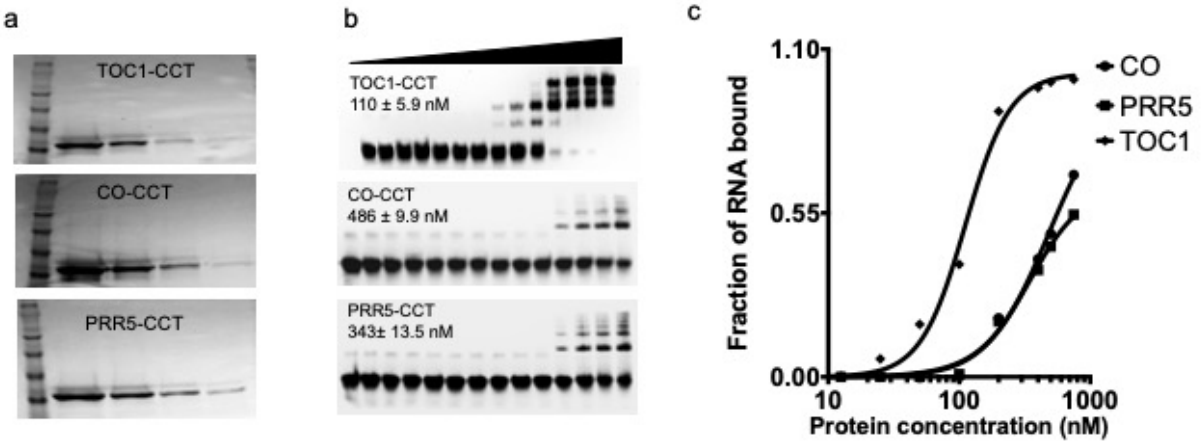


**Supplementary Fig. 6**

GST-tagged TOC1/CO/PRR5-CCT binds to RNA. (**a**) Coomassie-stained gel of purified TOC1/CO/PRR5-CCT domain. (**b**) EMSA showing recombinant TOC1/CO/PRR5-CCT binds to DNA and RNA. (**c**) Complete binding curves for TOC1/CO/PRR5-CCT domain to a G-rich RNA motif. The data were fitted by using a single exponential Hill equation.

**Supplementary Fig. 7:** Gel filtration analysis of CCT and RNA binding.

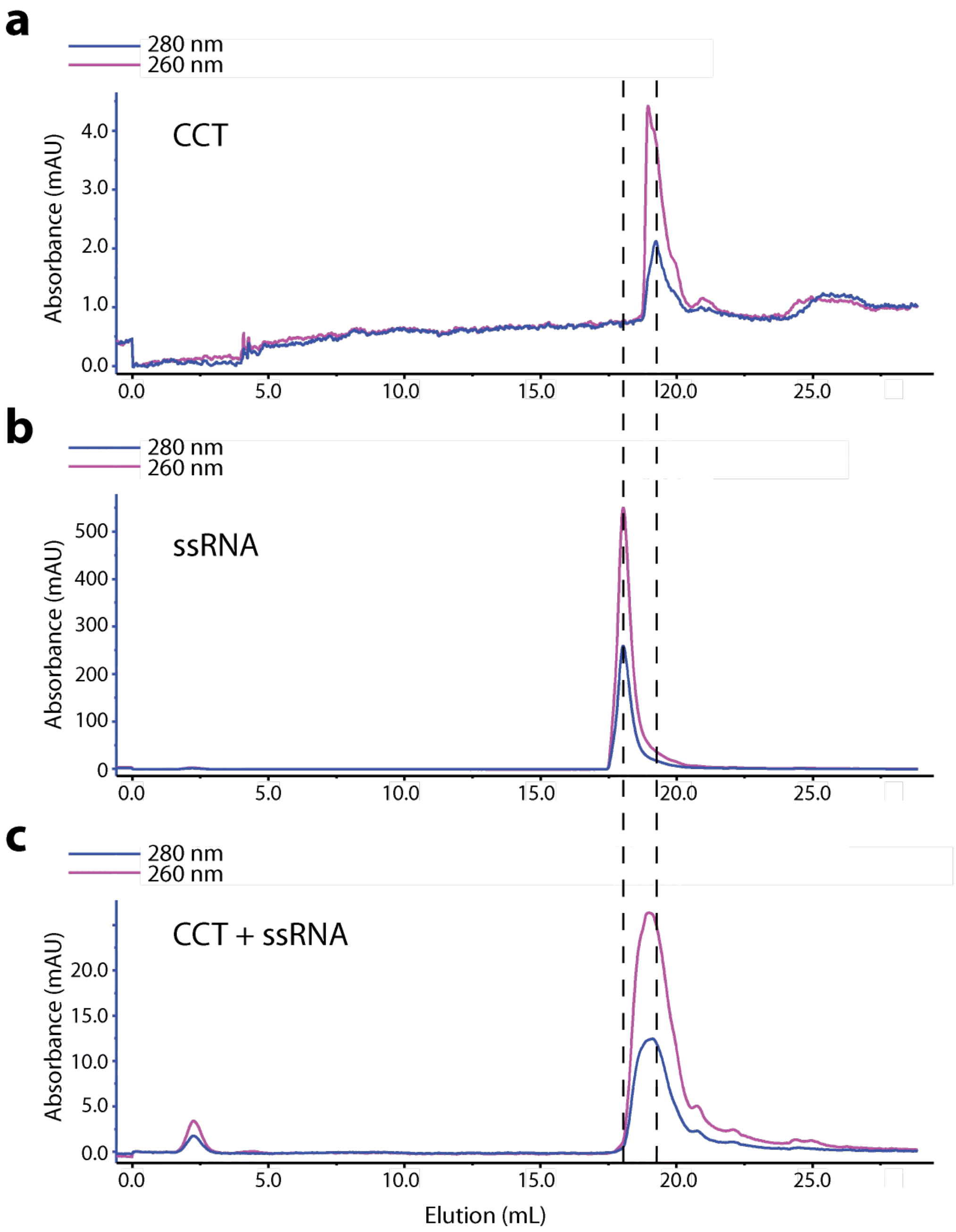


**Supplementary Fig. 7**

Gel filtration analysis of CCT and RNA binding. (a-c) Gel filtration profiles of free CCT (200 μM) (a), free RNA (5' UGG AGG GGA 3') (200 μM) (b), and mixture of CCT (200 μM) + RNA (200 μM). Samples (200 μL each) were loaded onto a Superose 6 Increase 10/300 GL column (bed volume: 24 mL) with a flow rate of 0.5 mL/min. The buffer for the runs was 10 mM Tris, 50 mM KCl, 5 mM MgCl2, 5 mM TCEP (Tris(2-carboxyethyl) phosphine), pH 7.0.

**Supplementary Fig. 8** Secondary structures of CCT predicated by PSIPRED.

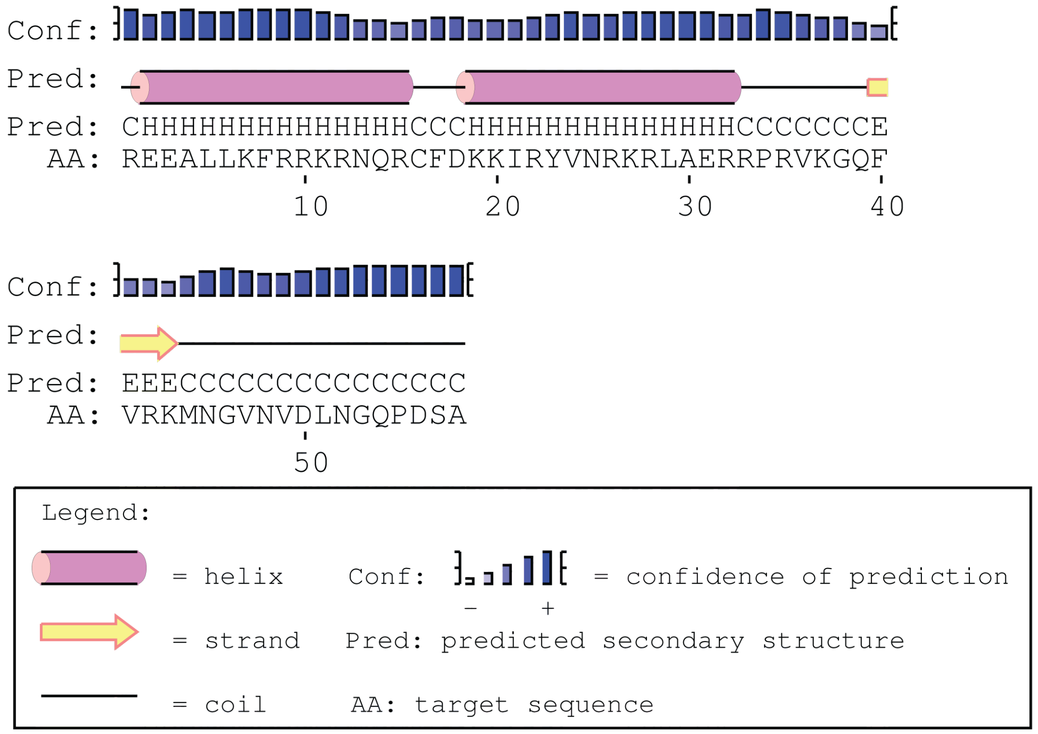


**Supplementary Fig. 8**

Secondary structures of CCT predicated by PSIPRED.

**Supplementary Fig. 9** Binding studies of CCT and RNA by NMR.

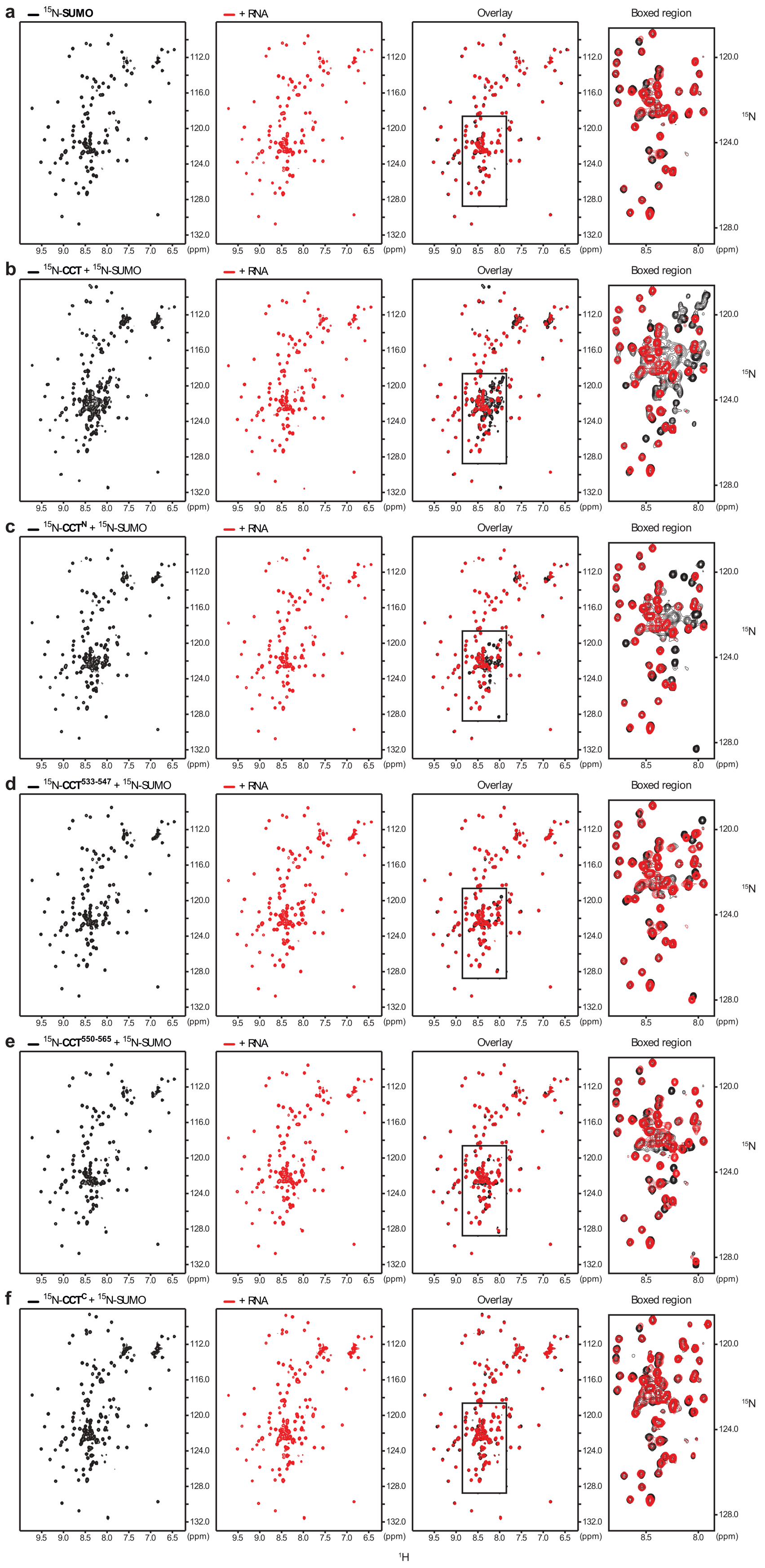

**Supplementary Fig. 9**

Binding studies of CCT and RNA by NMR.（**a**）$^{15}N,^{1}H$-HSQC spectra of free $^{15}N$-labeled SUMO ($1^{st}$ column); $^{15}N$-labeled SUMO mixed with unlabeled RNA (5' UGG AGG GGA 3') ($2^{nd}$ column); overlaid spectra ($3^{rd}$ column); zoomed-in view of the boxed region in the spectra in the $3^{rd}$ column ($4^{th}$ column).（**b-f**）Same as in (**a**) except for the additional presence of $^{15}N$-labeled CCT constructs ((**b**) CCT, (**c**) $CCT^{N}$, (**d**) $CCT^{533\text{-}547}$, (**e**) $CCT^{550\text{-}565}$, and (**f**) $CCT^{C}$).

**Supplementary Fig. 10** Backbone and sidechain assignment of $^{15}N$,$^{13}C$-labeled

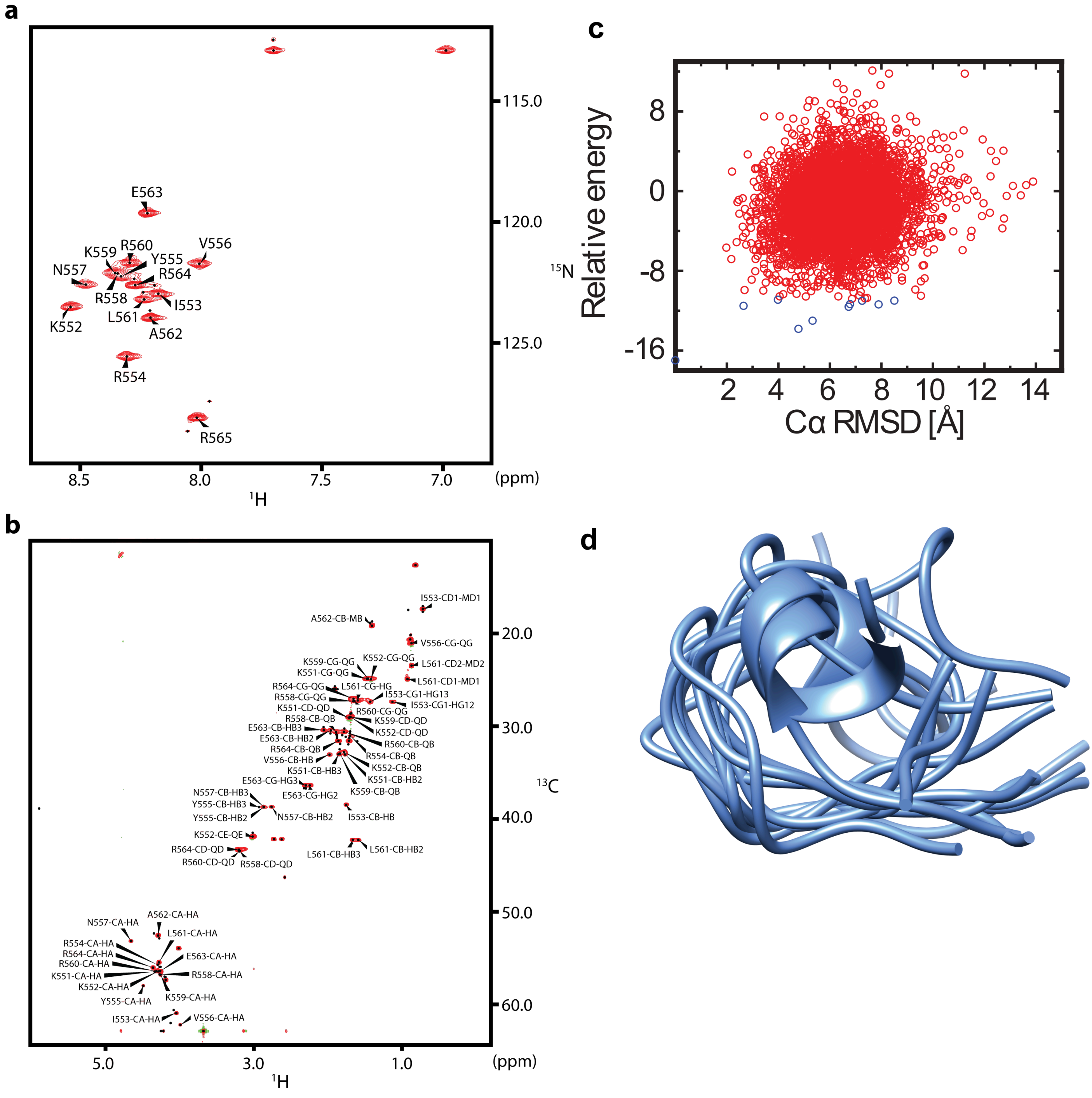


CCT$^{550-565}$ bound to RNA.

**Supplementary Fig. 10**

Backbone and sidechain assignment of $^{15}N,^{13}C$-labeled $CCT^{550-565}$ bound to RNA. (**a**) $^{15}N,^{1}H$-HSQC spectrum of $^{15}N,^{13}C$-labeled $CCT^{550-565}$ in complex with unlabeled RNA (5' UGG AGG GGA 3'). (**b**) $^{13}C,^{1}H$-HSQC spectrum of the same $CCT^{550-565}$-RNA complex in (**a**). Peaks in both spectra are labeled with assigned residues. (**c**) Plot of CS-Rosetta relative energy versus Cα RMSD of 5000 chemical shift guided structural models of $CCT^{550-565}$ bound to RNA. (**d**) Superposition of the ten lowest energy structural models.

**Supplementary Fig. 11** Mutational analysis of TOC1-CCT domain and RNA binding.

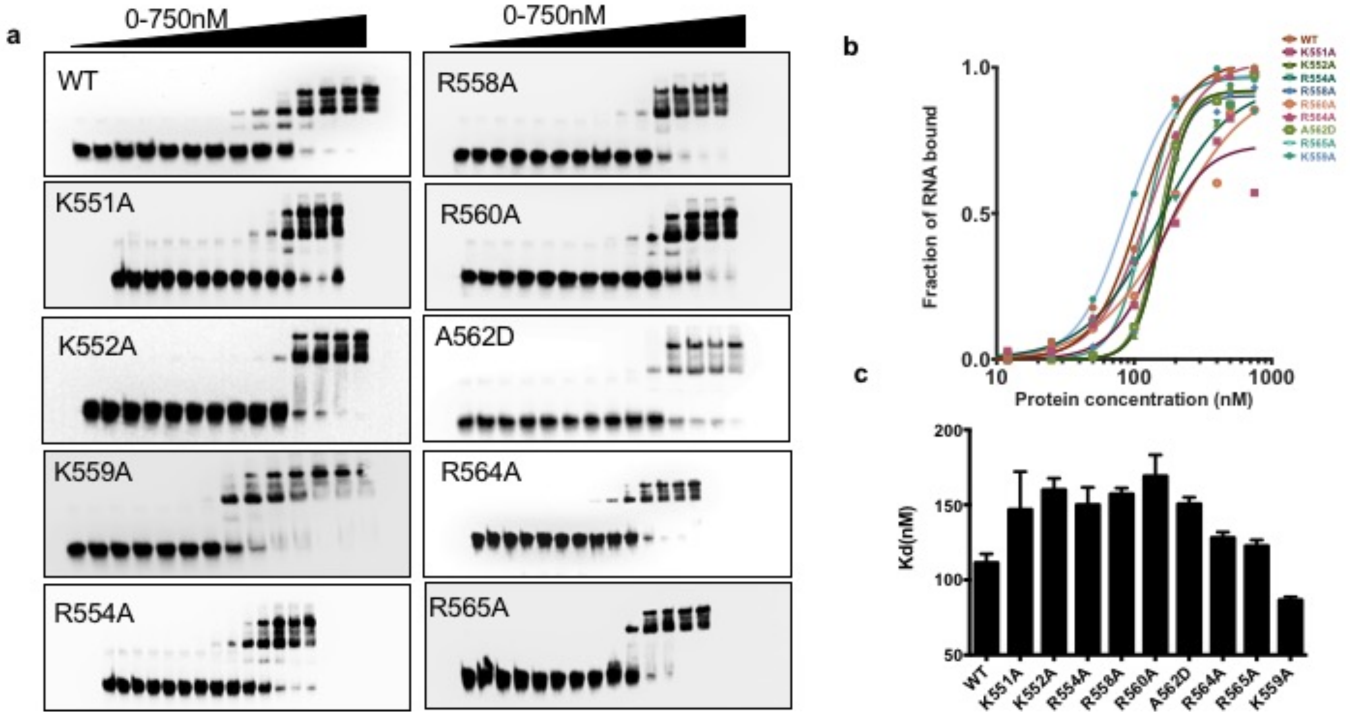


**Supplementary Fig. 11**

Mutational analysis of TOC1-CCT domain and RNA binding. (**a**) EMSA experiments using TOC1-CCT mutants (K551A, K552A, R554A, R558A, K559A, R560A, A562D, R564A, R565A). (**b**) Binding curves of TOC1-CCT mutants. (**c**) Binding affinities between TOC1-CCT mutants and G-rich RNA motif.

**Supplementary Fig. 12** Functional analysis of TOC1-CCT domain mutations *in vivo*.

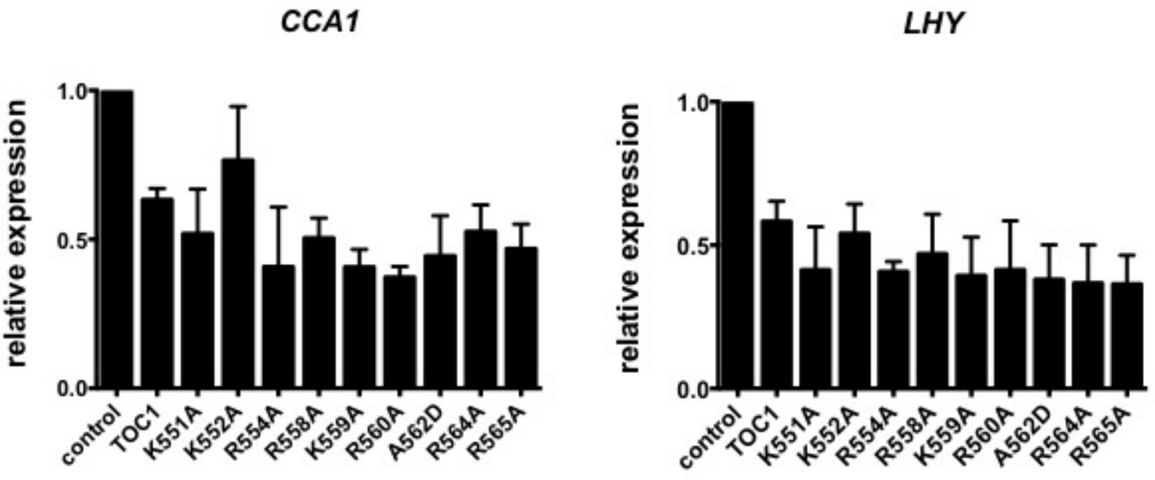


**Supplementary Fig. 12**

Functional analysis of TOC1-CCT domain mutations *in vivo*. Relative expression of TOC1 targets CCA1 and LHY in protoplast, overexpressing TOC1 and different TOC1 mutants. Statistical significance was calculated by one-way ANOVA (*$P < 0.05$).

**Supplementary Table S1: NMR sample conditions.**

| Sample | Experiment(s) | Sample details | Remark |
|---|---|---|---|
| $^{15}$N-SUMO<br>+<br>Ulp1 | 2D<br>$^{15}$N, $^{1}$H-HSQC | Concentration: 100 μM : 1 μM<br>Buffer: 10 mM Tris, 50 mM KCl, 5 mM $MgCl_2$, 5 mM TCEP, pH 7.0, 20 μM DSS, 95% $H_2O$/5% $D_2O$<br>Volume: 400 μL<br>Tube: Bruker shaped tube | 25 °C<br>Fig. 3b and Supplementary Fig. S9a |
| $^{15}$N-SUMO<br>+<br>RNA<br>+<br>Ulp1 | 2D<br>$^{15}$N, $^{1}$H-HSQC | Concentration: 100 μM : 200 μM : 1 μM<br>Buffer: 10 mM Tris, 50 mM KCl, 5 mM $MgCl_2$, 5 mM TCEP, pH 7.0, 20 μM DSS, 95% $H_2O$/5% $D_2O$<br>Volume: 400 μL<br>Tube: Bruker shaped tube | 25 °C<br>Fig. 3b and Supplementary Fig. S9a |
| $^{15}$N-CCT<br>(or its fragments: $CCT^{N}$, $CCT^{533\text{-}547}$, $CCT^{550\text{-}565}$, $CCT^{C}$)<br>+<br>$^{15}$N-SUMO<br>+<br>Ulp1 | 2D<br>$^{15}$N, $^{1}$H-HSQC | Concentration: 100 μM : 100 μM : 1 μM<br>Buffer: 10 mM Tris, 50 mM KCl, 5 mM $MgCl_2$, 5 mM TCEP, pH 7.0, 20 μM DSS, 95% $H_2O$/5% $D_2O$<br>Volume: 400 μL<br>Tube: Bruker shaped tube | 25 °C<br>Fig. 3b and Supplementary Fig. S9b to S9f |
| $^{15}$N-CCT<br>(or its fragments: $CCT^{N}$, $CCT^{533\text{-}547}$, $CCT^{550\text{-}565}$, $CCT^{C}$)<br>+<br>$^{15}$N-SUMO<br>+<br>RNA<br>+<br>Ulp1 | 2D<br>$^{15}$N, $^{1}$H-HSQC | Concentration: 100 μM : 100 μM : 200 μM : 1 μM<br>Buffer: 10 mM Tris, 50 mM KCl, 5 mM $MgCl_2$, 5 mM TCEP, pH 7.0, 20 μM DSS, 95% $H_2O$/5% $D_2O$<br>Volume: 400 μL<br>Tube: Bruker shaped tube | 25 °C<br>Fig. 3b and Supplementary Fig. S9b to S9f |
| $^{15}$N,$^{13}$C-$CCT^{550\text{-}565}$<br>+<br>$^{15}$N,$^{13}$C-SUMO<br>+<br>RNA | 3D<br>$^{13}$C-edited, $^{12}$C-filtered NOESY | Concentration: 0.5 mM : 0.1 mM : 2 mM<br>Buffer: 10 mM Tris, 50 mM KCl, 5 mM $MgCl_2$, pH 7.0, 20 μM DSS, 99.8% D2O<br>Volume: 400 μL<br>Tube: Bruker shaped tube | 25 °C<br>Fig. 3c |
| $^{15}$N,$^{13}$C-$CCT^{550\text{-}565}$<br>+<br>RNA | 3D<br>$^{13}$C-edited, $^{12}$C-filtered NOESY | Concentration: 0.85 mM : 1.7 mM<br>Buffer: 10 mM Tris, 50 mM KCl, 5 mM $MgCl_2$, pH 7.0, 20 μM DSS, 0.02% $NaN_3$, 99.96% D2O<br>Volume: 400 μL<br>Tube: Bruker shaped tube | 25 °C<br>Fig. 3c |
| $^{15}$N,$^{13}$C-$CCT^{550\text{-}565}$<br>+<br>RNA | 3D<br>HNCACB, HNCOCACB, HNCACO, HNCO, HBHACONH, HCCCONH, CCCONH<br>2D<br>$^{15}$N, $^{1}$H-HSQC<br>$^{13}$C, $^{1}$H-HSQC | Concentration: 0.85 mM : 1.7 mM<br>Buffer: 10 mM Tris, 50 mM KCl, 5 mM $MgCl_2$, pH 7.0, 20 μM DSS, 0.02% $NaN_3$, 95% $H_2O$/5% $D_2O$<br>Volume: 400 μL<br>Tube: Bruker shaped tube | 25 °C<br>Supplementary Fig. S10 |

**Supplementary Table S2: List of parameters for 2D HSQC experiments.**

| Pulse program (Experiment) | Parameters | *t1* | *t2* |
|---|---|---|---|
| | | $^{15}$N (or $^{13}$C) | $^{1}$H |
| hsqcfpf3fpphwg3 ($^{15}$N,$^{1}$H-HSQC) | Complex points | 256 | 1160 |
| | Acquisition (s) | 0.0724 | 0.0689 |
| | Sweep width (ppm) | 29.0000 | 14.0027 |
| hsqcetfpf3gpsi2 ($^{15}$N,$^{1}$H-HSQC) | Complex points | 254 | 980 |
| | Acquisition (s) | 0.0802 | 0.0679 |
| | Sweep width (ppm) | 26.0000 | 11.9966 |
| hsqcctetgpsp ($^{13}$C,$^{1}$H-HSQC) (aliased in $^{13}$C dimension) | Complex points | 302 | 816 |
| | Acquisition (s) | 0.0290 | 0.0679 |
| | Sweep width (ppm) | 34.50000 | 9.9972 |
| Hsqcctetgpsp ($^{13}$C,$^{1}$H-HSQC) | Complex points | 302 | 816 |
| | Acquisition (s) | 0.0143 | 0.0679 |
| | Sweep width (ppm) | 70.0000 | 9.9972 |

**Supplementary Table S3: List of parameters for 3D assignment and NOE experiments.** The two NOESY experiments listed below were performed on two different D2O samples [$^{15}$N,$^{13}$C-CCT$^{550\text{-}565}$ + $^{15}$N,$^{13}$C-SUMO + RNA; $^{15}$N,$^{13}$C-CCT$^{550\text{-}565}$ + RNA], respectively. All other experiments were performed on the H2O sample [$^{15}$N,$^{13}$C-CCT$^{550\text{-}565}$ + RNA]. Refer to Table S1 for sample details.

| **Pulse program (Experiment)** | **Parameters** | ***t1*** | ***t2*** | ***t3*** |
|---|---|---|---|---|
| | | **$^{1}$H (or $^{13}$C)** | **$^{15}$N** | **$^{1}$H** |
| noesyhsqcgpwgx13d ($^{13}$C-edited, $^{12}$C-filtered NOESY) (sweep width: 20.0000 ppm) | Complex points | 144 | 68 | 864 |
| | Acquisition (s) | 0.0200 | 0.0112 | 0.0599 |
| | Sweep width (ppm) | 6.0000 | 20.0000 | 11.9966 |
| noesyhsqcgpwgx13d ($^{13}$C-edited, $^{12}$C-filtered NOESY) (sweep width: 34.5000 ppm) | Complex points | 144 | 68 | 864 |
| | Acquisition (s) | 0.0200 | 0.0065 | 0.0599 |
| | Sweep width (ppm) | 6.0000 | 34.5000 | 11.9966 |
| hncacbgp3d (HNCACB) | Complex points | 132 | 76 | 980 |
| | Acquisition (s) | 0.0069 | 0.0240 | 0.0679 |
| | Sweep width (ppm) | 62.8470 | 26.0000 | 11.9966 |
| hncocacbgp3d (HNCOCACB) | Complex points | 132 | 74 | 980 |
| | Acquisition (s) | 0.0069 | 0.0234 | 0.0679 |
| | Sweep width (ppm) | 62.8470 | 26.0000 | 11.9966 |
| hncacogp3d (HNCACO) | Complex points | 90 | 74 | 980 |
| | Acquisition (s) | 0.0300 | 0.0234 | 0.0679 |
| | Sweep width (ppm) | 9.9219 | 26.0000 | 11.9966 |
| hncogp3d (HNCO) | Complex points | 90 | 74 | 980 |
| | Acquisition (s) | 0.0300 | 0.0234 | 0.0679 |
| | Sweep width (ppm) | 9.9219 | 26.0000 | 11.9966 |
| hbhaconhgp3d (HBHACONH) | Complex points | 104 | 76 | 980 |
| | Acquisition (s) | 0.0130 | 0.0240 | 0.0679 |
| | Sweep width (ppm) | 6.6541 | 26.0000 | 11.9966 |
| hccconhgpwg3d2 (HCCCONH) | Complex points | 104 | 76 | 980 |
| | Acquisition (s) | 0.0130 | 0.0240 | 0.0679 |
| | Sweep width (ppm) | 6.6541 | 26.0000 | 11.9966 |
| hccconhgpwg3d3 (CCCONH) | Complex points | 156 | 76 | 980 |
| | Acquisition (s) | 0.0081 | 0.0240 | 0.0679 |
| | Sweep width (ppm) | 64.0000 | 26.0000 | 11.9966 |
